# Unveiling the Robustness of Machine Learning Models in Classifying COVID-19 Spike Sequences

**DOI:** 10.1101/2023.08.24.554651

**Authors:** Sarwan Ali, Pin-Yu Chen, Murray Patterson

**Author notes:** {, }.

## Abstract

In the midst of the global COVID-19 pandemic, a wealth of data has become available to researchers, presenting a unique opportunity to investigate the behavior of the virus. This research aims to facilitate the design of efficient vaccinations and proactive measures to prevent future pandemics through the utilization of machine learning (ML) models for decision-making processes. Consequently, ensuring the reliability of ML predictions in these critical and rapidly evolving scenarios is of utmost importance. Notably, studies focusing on the genomic sequences of individuals infected with the coronavirus have revealed that the majority of variations occur within a specific region known as the spike (or S) protein. Previous research has explored the analysis of spike proteins using various ML techniques, including classification and clustering of variants. However, it is imperative to acknowledge the possibility of errors in spike proteins, which could lead to misleading outcomes and misguide decision-making authorities. Hence, a comprehensive examination of the robustness of ML and deep learning models in classifying spike sequences is essential. In this paper, we propose a framework for evaluating and benchmarking the robustness of diverse ML methods in spike sequence classification. Through extensive evaluation of a wide range of ML algorithms, ranging from classical methods like naive Bayes and logistic regression to advanced approaches such as deep neural networks, our research demonstrates that utilizing *k*-mers for creating the feature vector representation of spike proteins is more effective than traditional one-hot encoding-based embedding methods. Additionally, our findings indicate that deep neural networks exhibit superior accuracy and robustness compared to non-deep-learning baselines. To the best of our knowledge, this study is the first to benchmark the accuracy and robustness of machine-learning classification models against various types of random corruptions in COVID-19 spike protein sequences. The benchmarking framework established in this research holds the potential to assist future researchers in gaining a deeper understanding of the behavior of the coronavirus, enabling the implementation of proactive measures and the prevention of similar pandemics in the future.

## 1 Introduction

In January 2020, a new RNA coronavirus was discovered, marking the onset of the ongoing COVID-19 pandemic. Through the utilization of sequencing technology and phylogenetic analysis, the scientific community determined that this novel coronavirus shares a 50% similarity with the Middle-Eastern Respiratory Syndrome Coronavirus (MERS-CoV), a 79% sequencing similarity with the Severe Acute Respiratory Syndrome Coronavirus (SARS-CoV), commonly known as “SARS,” and over 85% similarity with coronaviruses found in bats. Subsequent research confirmed bats as the probable reservoir of these coronaviruses; however, due to the ecological separation between bats and humans, it is believed that other organisms might have served as intermediate hosts. Based on comprehensive scientific evidence, the International Committee on Taxonomy of Viruses officially designated the novel RNA virus as SARS-CoV-2 [31,26,32].

RNA viruses commonly introduce errors during the replication process, resulting in mutations that become part of the viral genome after multiple replications within a single host. This leads to the formation of a diverse population of viral quasispecies. However, SARS-CoV-2 possesses a highly effective proofreading mechanism facilitated by a nonstructural protein 14 (nsp14), resulting in a mutation rate approximately ten times lower than that of typical RNA viruses. On average, it is estimated that SARS-CoV-2 acquires 33 genomic mutations per year. Some of these mutations confer advantages, leading to the emergence of more infectious variants of SARS-CoV-2. These variants can be distinguished by a small number of specific mutations, and the relatively slow accumulation of mutations makes it unlikely for minor sequence perturbations or errors to cause confusion between different variants. Additionally, most of these mutations occur in the S gene, which encodes the spike protein responsible for the surface characteristics of the virus. Consequently, characterizing variants based on the spike proteins transcribed from the genome is sufficient for the classification task [25,28,21].

The decreasing cost of next-generation sequencing (NGS) technology has significantly facilitated SARS-CoV-2 whole-genome sequencing (WGS) by researchers worldwide. The Centers for Disease Control and Prevention (CDC) in the United States have provided comprehensive resources, tools, and protocols for SARS-CoV-2 WGS using various sequencing platforms such as Illumina, PacBio, and Ion Torrent. Additionally, the Global Initiative on Sharing All Influenza Data (GISAID) hosts the largest SARS-CoV-2 genome sequencing dataset to date, encompassing millions of sequences. This unparalleled volume of genomic data generation and its easy accessibility have enabled researchers to delve into the molecular mechanisms, genetic variability, evolutionary progression, and the potential for the emergence and spread of novel virus variants. However, the sheer magnitude of this data surpasses the capabilities of existing methods like Nextstrain [16] and even the more recent IQTREE2 [24] by several orders of magnitude, presenting a significant Big Data challenge. Consequently, alternative approaches focusing on clustering and classification of sequences have emerged in recent literature [23,2,5,3], demonstrating promising accuracy and scalability properties. These methods offer viable solutions for identifying major variants and addressing the challenges associated with the extensive volume of genomic data.

However, several challenges persist in studying the evolutionary and transmission patterns of SARS-CoV-2 [13,6] and other viruses. One of these challenges arises from sequencing errors, which can be mistakenly identified as mutations during analysis. These errors can result from various sources, including contamination during sample preparation, sequencing technology limitations, or genome assembly methodologies. To address this issue, computational biologists typically employ filtering techniques to remove sequences with errors or mask specific sequence fragments that exhibit errors. For instance, in the case of GISAID [14] sequences, each sequence represents a consensus sequence derived from the viral population within an infected individual. This consensus sequence averages out minor variations present in the population, providing a representative snapshot of the SARS-CoV-2 variant carried by the patient. Although this consensus sequence accurately captures the predominant variant, it comes at the cost of losing valuable information about these minor variations. However, given enough time and within an immunocompromised individual, these minor variations can undergo significant evolution and become dominant, as observed in the emergence of the Alpha variant [12].

What this means is that many machine learning approaches towards clustering and classification of sequences [2,5,3] have been operating under rather idealized conditions of virtually error-free consensus sequences. Moreover, these methods rely on a *k*-mer based feature vector representation — an approach that does not even rely on the alignment of the sequences, something which can also introduce bias [15]. Such a framework should easily cope with errors as well — something machine learning approaches can do very naturally [11]. There is hence a great need for some way to reliably benchmark such methods for robustness to errors, which is what we carry out in this paper.

We highlight the main contributions of this paper as follows:

– We propose several ways of introducing errors into spike sequences which reflect the error profiles of modern NGS technologies such as Illumina and PacBio;

– We demonstrate that the *k*-mer based approach is more robust to such errors when compared to the baseline (one-hot encoding); and

– We show that deep learning is generally more robust in handling these errors than machine learning models.

Moreover, we extend our error testing procedure as a framework for benchmarking the performance of different ML methods in terms of classification accuracy and robustness to different types of simulated random errors in the sequences. The two types of errors that we introduce are “consecutive” and “random” errors (see Sec. 3.4). Random errors are just point mutations, which happen uniformly at random along the protein sequence, simulating closely the behavior of Illumina sequencing technolgies [30]. Consecutive errors on the other hand, are small subsequences of consecutive errors, which can model insertion-deletion (indel) errors which are common in third generation long-read technologies such as Pacific Biosciences (PacBio) SMRT sequencing [10].

This paper is structured as follows. In Sec. 2, we discuss related work. In Sec. 3 we discuss some approaches we benchmark, and then how we benchmark: the type of adversarial attacks we use. Sce. 4 details the experiments, and Sec. 5 gives the results. Finally, we conclude this paper in Sec. 6.

## 2 Related Work

The evaluation and benchmarking of the robustness of machine learning (ML) and deep learning (DL) approaches through adversarial attacks have gained popularity in the field of image classification [17]. However, there are related works that focus more specifically on molecular data. For instance, in [29], the authors present a set of realistic adversarial attacks to assess methods that predict chemical properties based on atomistic simulations, such as molecular conformation, reactions, and phase transitions. Additionally, in the context of protein sequences, the authors of [18] demonstrate that deep neural network-based methods like AlphaFold [19] and RoseTTAFold [7], which predict protein conformation, lack robustness. These methods produce significantly different protein structures when subjected to small, biologically meaningful perturbations in the protein sequence. Although our approach shares similarities with these works, our goal is classification. Specifically, we investigate how a small number of point mutations, simulating errors introduced by certain types of nextgeneration sequencing (NGS) technologies, can impact the downstream classification performed by various ML and DL approaches.

When it comes to obtaining numerical representations, a common approach involves constructing a kernel matrix and using it as input for traditional machine learning classifiers like support vector machines (SVM)[22,20]. However, these methods can be computationally expensive in terms of space complexity. In related works[3,21], efficient embedding methods for spike sequence classification and clustering are proposed. Nevertheless, these approaches either lack scalability or exhibit poor performance when dealing with larger datasets. Although some effort has been made to benchmark the robustness of machine learning models using genome (nucleotide) sequences [4], no such study has been conducted on the (spike) protein sequences (to the best of our knowledge).

## 3 Proposed Approach

In this section, we start by explaining the baseline model for spike sequence classification. After that, we will explain our deep learning model in detail.

### 3.1 One-Hot Encoding (OHE) Based Embedding

Authors in [21] propose that classification of viral hosts of the coronavirus can be done by using spike sequences only. For this purpose, a fixed-length onehot encoding-based feature vector is generated for the spike sequences. In the spike sequence, we have 21 unique characters (amino acids) that are *“ACDEFGHIKLMNPQRSTVWXY”*. Also, note that the length of each spike sequence is 1273 with the stopping character ‘*’ at the 1274^*th*^ position. When we design the OHE-based numerical vector for the spike sequence, the length of the numerical vector will be 21 × 1273 = 26733. This high dimensionality could create the problem of “Curse of Dimensionality (CoD)”. To solve the CoD problem, any dimensionality reduction method can be used, such as Principal Component Analysis [1]. After reducing the dimensions of the feature vectors, classical Machine Learning (ML) algorithms can be applied to classify the spike sequences. One major problem with such OHE-based representation is that it does not preserve the order of the amino acids very efficiently [2]. If we compute the pair-wise Euclidean distance between any two OHE-based vectors, the overall distance will not be affected if a random pair of amino acids are swapped for those two feature vectors. Since the order of amino acids is important in the case of sequential data, OHE fails to give us efficient results [2]. In this paper, we use OHE as a baseline embedding method.

### 3.2 *k*-mers Based Representation

A popular approach to preserve the ordering of the sequential information is to take the sliding window-based substrings (called mers) of length *k*. This *k*mers-based representation is recently proven to be useful in classifying the spike sequences effectively [2]. In this approach, first, the *k*-mers of length *k* are computed for each spike sequence. Then a fixed length frequency vector is generated corresponding to each spike sequence, which contains the count of each *k*-mer in that sequence. One advantage of using *k*-mers based approach is that it is an *“alignment-free”* method, unlike OHE, which requires the sequences to be aligned to the reference genome.

#### Remark 1

Sequence alignment is an expensive process and requires reference sequence (genome) [8,9]. It may also introduce bias into the result [15].

The total number of *k*-mers in a given spike sequence are:

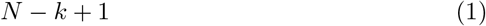

where *N* is the length of the sequence. The variable *k* is the user-defined parameter. In this paper, we take *k* = 3 (decided empirically). Since we have 1273 length spike sequences, the total number of *k*-mers that we can have for any spike sequence is 1273 – 3 + 1 = 1271.

### Frequency Vector Generation

After generating the *k*-mers, the next step is to generate the fixed-length numerical representation (frequency vector) for the set of *k*-mers in a spike sequence. Let the set of amino acids in the whole dataset is represented by alphabet *Σ*. Now, length of the frequency vector will be | *Σ* | ^*k*^ (all possible combinations of *k*-mers in *Σ* of length *k*). Recall that in our dataset, we have 21 unique amino acids in any spike sequence. Therefore, the length of frequency vector in our case would be 21^3^ = 9261 (when we take *k* = 3).

Note that CoD could be a problem in the case of *k*-mers based numerical representation of the spike sequence. To deal with this problem, authors in [3] use an approximate kernel method that map such vectors into low dimensional euclidean space using an approach, called Random Fourier Features (RFF) [27]. Unlike kernel trick, which compute the inner product between the lifted data points *ϕ* (i.e. ⟨*ϕ*(*a*), *ϕ*(*b*)⟩ = *f* (*a, b*), where *a, b* ∈ ℜ^*d*^ and f(a,b) is any positive definite function), RFF maps the data into low dimensional euclidean inner product space. More formally:

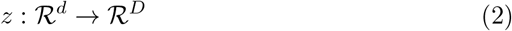

RFF tries to approximate the inner product between any pair of transformed points.

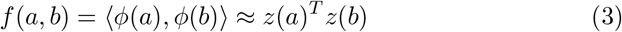

where *z* is low dimensional representation. Since *z* is the approximate representation, we can use it as an input for the classical ML models and analyse their behavior (as done in Spike2Vec [3]). However, we show that such approach performs poorly on larger size datasets (hence poor scalability).

### 3.3 Keras Classifier

We use a deep learning-based model called the Keras Classification model (also called Keras classifier) to further improve the performance that we got from Spike2Vec. For keras classifier, we use a sequential constructor. It contains a fully connected network with one hidden layer that contains neurons equals to the length of the feature vector (i.e. 9261). We use “rectifier” activation function for this classifier. Moreover, we use “softmax” activation function in the output layer. At last, we use the efficient Adam gradient descent optimization algorithm with “sparse categorical crossentropy” loss function (used for multi-class classification problem). It computes the crossentropy loss between the labels and predictions. The batch size and number of epochs are taken as 100 and 10, respectively for training the model. For the input to this keras classifier, we separately use OHE and *k*-mers based frequency vectors.

#### Remark 2

Note that we are using “sparse categorical crossentropy” rather than simple “categorical crossentropy” because we are using integer labels rather than one-hot representation of labels.

### 3.4 Adversarial Examples Creation

We use two types of approaches to generate adversarial examples so that we can test the robustness of our proposed model. These approaches are “Random Error generation” and “Consecutive Error generation”.

In random error generation, we randomly select a fraction of spike sequences (we call them the set of errored sequences for reference) from the test set (i.e. 5%, 10%, 15%, and 20%). For each of the spike sequence in the set of errored sequences, we randomly select a fraction of amino acids (i.e. 5%, 10%, 15%, and 20%) and flip their value randomly. At the end, we replace these errored sequences set with the corresponding set of original spike sequences in the test set. The ideas is that this simulates the errors made by NGS technologies such as Illumina [30].

In consecutive error generation, the first step is the same as in random error generation (getting random set of spike sequences from the test set “set of errored sequences”). For this set of errored sequences, rather than randomly flipping a specific percentage of amino acid’s values for each spike sequence (i.e. 5%, 10%, 15%, and 20%), we flip the values for the same fraction of amino acids but those amino acids are consecutive and at random position in the spike sequence. More formally, it is a consecutive set of amino acids (at random position) in the spike sequence for which we flip the values. At the end, we replace these errored sequences set with the corresponding set of original spike sequences in the test set. The idea is that this simulates indel errors, which are frequencly found in third generation long-read technologies such as PacBio [10].

Using the two approaches to generate adversarial examples, we generate a new test set and evaluate the performance of the ML and deep learning models. To measure the performance, we also apply two different strategies. One strategy is called Accuracy and the other is called robustness. In the case of the Accuracy, we compute the average accuracy, precision, recall, F1 (weighted), F1 (Macro), and ROC-AUC for the whole test set including adversarial and non-adversarial examples. For our second strategy (robustness), we only consider the adversarial examples (set of errored spike sequences) rather than considering the whole test set and compute average accuracy, precision, recall, F1 (weighted), F1 (Macro), and ROC-AUC for them.

## 4 Experimental Setup

All experiments are conducted using an Intel(R) Xeon(R) CPU E7-4850 v4 @ 2.10GHz having Ubuntu 64 bit OS (16.04.7 LTS Xenial Xerus) with 3023 GB memory. Our pre-processed data is also available online^3^, which can be used after agreeing to terms and conditions of GISAID^4^. For the classification algorithms, we use 10% data for training and 90% for testing. Note that our data split and pre-processing follow those of [3].

### 4.1 Dataset Statistics

We used the (aligned) spike protein from a popular and publicly available database of SARS-CoV-2 sequences, GISAID. In our dataset, we have 2, 519, 386 spike sequences along with the COVID-19 variant information. The total number of unique variants in our dataset are 1327. Since not all variants have significant representation in our data, we only select the variants having more than 10, 000 sequences. After this preprocessing, we are left with 1, 995, 195 spike sequences [3].

## 5 Results and Discussion

In this section, we first show the comparison of our deep learning model with the baselines. We then show the results for the two approaches for adversarial examples generation and compare different ML and DL methods. Overall, we elucidate our key findings in the following subsections.

### 5.1 Effectiveness of Deep Learning

Table 1 contains the (accuracy) results for our keras classifier and its comparison with different ML models namely Naive Nayes (NB), Logistic Regression (LR), and Ridge Classifier (RC). For keras classifier, we use both OHE and *k*-mers-based embedding approaches separately. We can observe from the results that keras classifier with *k*-mers based frequency vectors is by far the best approach as compared to the other baselines.

**Table 1:** Variants Classification Results (10% training set and 90% testing set) for top 22 variants (1995195 spike sequences).

| Approach | Embed. Method | ML Algo. | Acc. | Prec. | Recall | F1 weigh. | F1 Macro | ROC AUC | Train. run-time (sec.) |
| --- | --- | --- | --- | --- | --- | --- | --- | --- | --- |
| ML Algo. | OHE | NB | 0.30 | 0.58 | 0.30 | 0.38 | 0.18 | 0.59 | 1164.5 |
|  |  | LR | 0.57 | 0.50 | 0.57 | 0.49 | 0.19 | 0.57 | 1907.5 |
|  |  | RC | 0.56 | 0.48 | 0.56 | 0.48 | 0.17 | 0.56 | 709.2 |
|  | Spike2Vec | NB | 0.42 | 0.79 | 0.42 | 0.52 | 0.39 | 0.68 | 1056.0 |
|  |  | LR | 0.68 | 0.69 | 0.68 | 0.65 | 0.49 | 0.69 | 1429.1 |
|  |  | RC | 0.67 | 0.68 | 0.67 | 0.63 | 0.44 | 0.67 | 694.2 |
| Deep Learning | One-Hot | Keras Classifier | 0.61 | 0.58 | 0.61 | 0.56 | 0.24 | 0.61 | 28971.5 |
| | $k$-mers | Keras Classifier | **0.87** | **0.88** | **0.87** | **0.86** | **0.71** | **0.85** | 13296.2 |

To test the robustness of these ML and DL methods, we use both “consecutive error generation” and “random error generation” separately. Table 2 shows the (accuracy) results (using keras classifier with *k*-mers because that was the best model from Table 1) for the consecutive error generation method (using different fraction of spike sequences from the test set and different fraction of amino acids flips in each spike sequence). We can observe that keras classifier is able to perform efficiently even with higher proportion of error.

**Table 2:**
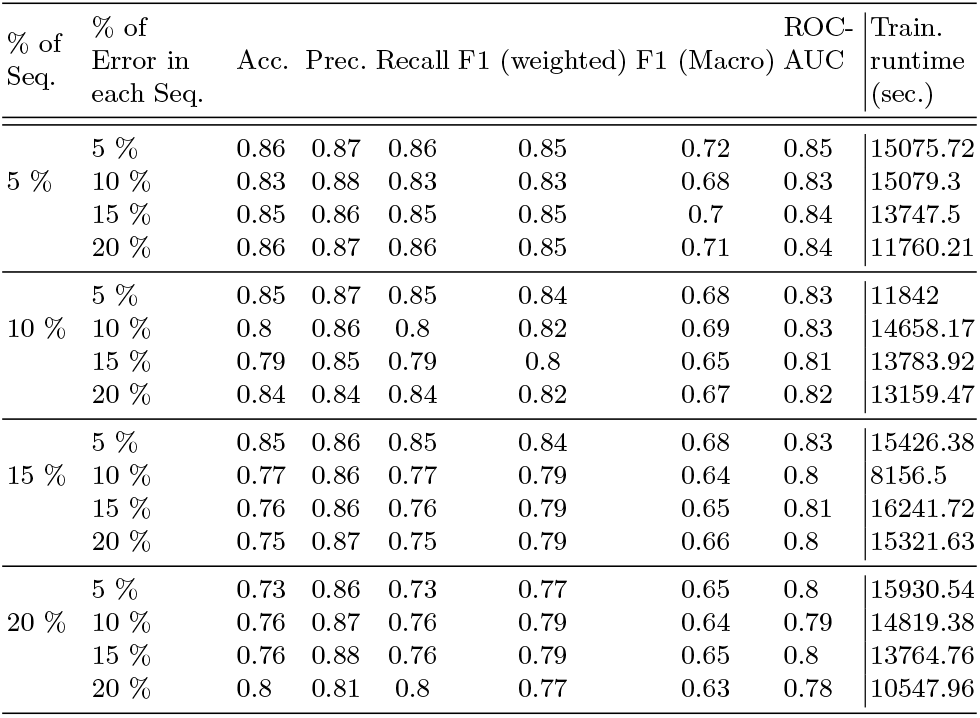
Accuracy results for the whole test set (consecutive error seq.) for Keras Classifier with *k*-mers and different % of errors.

The robustness results for the consecutive error generation method are given in Table 3. Although we cannot see any clear pattern in this case, the keras classifier is giving us comparatively higher performance in some of the settings.

**Table 3:** Robustness (only error seq. in the test set) results (consecutive error seq.) for Keras Classifier with *k*-mers and different % of errors.

| % of Seq. | % of Error in each Seq. | Acc. | Prec. | Recall | F1 (weighted) | F1 (Macro) | ROC-AUC | Train. runtime (sec.) |
| --- | --- | --- | --- | --- | --- | --- | --- | --- |
| 5 % | 5 % | 0.6 | 0.57 | 0.6 | 0.54 | 0.1 | 0.55 | 13882.03 |
|  | 10 % | 0.15 | 0.62 | 0.15 | 0.1 | 0.04 | 0.52 | 8297.78 |
|  | 15 % | 0.12 | 0.51 | 0.12 | 0.03 | 0.01 | 0.5 | 11943.5 |
|  | 20 % | 0.6 | 0.44 | 0.6 | 0.48 | 0.07 | 0.53 | 13511.4 |
| 10 % | 5 % | 0.53 | 0.62 | 0.53 | 0.53 | 0.09 | 0.54 | 9475.31 |
|  | 10 % | 0.31 | 0.62 | 0.31 | 0.32 | 0.1 | 0.54 | 7137.31 |
|  | 15 % | 0.25 | 0.6 | 0.25 | 0.27 | 0.07 | 0.53 | 13399.18 |
|  | 20 % | 0.55 | 0.35 | 0.55 | 0.42 | 0.06 | 0.52 | 13232.93 |
| 15 % | 5 % | 0.43 | 0.76 | 0.43 | 0.46 | 0.24 | 0.61 | 13588.11 |
|  | 10 % | 0.54 | 0.58 | 0.54 | 0.54 | 0.11 | 0.56 | 14147.05 |
|  | 15 % | 0.06 | 0.63 | 0.06 | 0.02 | 0.02 | 0.51 | 13729.87 |
|  | 20 % | 0.06 | 0.64 | 0.06 | 0.03 | 0.02 | 0.5 | 13596.25 |
| 20 % | 5 % | 0.17 | 0.68 | 0.17 | 0.18 | 0.1 | 0.54 | 13503.11 |
|  | 10 % | 0.48 | 0.63 | 0.48 | 0.48 | 0.09 | 0.55 | 10777.34 |
|  | 15 % | 0.47 | 0.59 | 0.47 | 0.49 | 0.09 | 0.55 | 13550.35 |
|  | 20 % | 0.49 | 0.39 | 0.49 | 0.33 | 0.03 | 0.5 | 11960.26 |

Table 4 contains the accuracy results for the keras classifier (with *k*-mers based frequency vectors as input) with random errored sequences approach. We can observe that our DL model is able to maintain higher accuracy even with 20% of the spike sequences having some fraction of error in the test set. Similarly, the robustness results are given in Table 5.

**Table 4:** Accuracy results for the whole test set (random error seq.) for Keras Classifier with *k*-mers and different % of errors.

| % of Seq. | % of Error in each Seq. | Acc. | Prec. | Recall | F1 (weighted) | F1 (Macro) | ROC-AUC | Train. runtime (sec.) |
| --- | --- | --- | --- | --- | --- | --- | --- | --- |
| 5 % | 5 % | 0.85 | 0.88 | 0.85 | 0.84 | 0.69 | 0.84 | 9338.87 |
|  | 10 % | 0.84 | 0.87 | 0.84 | 0.85 | 0.72 | 0.85 | 14365.47 |
|  | 15 % | 0.84 | 0.87 | 0.84 | 0.83 | 0.70 | 0.84 | 14996.06 |
|  | 20 % | 0.82 | 0.84 | 0.82 | 0.81 | 0.68 | 0.83 | 10958.00 |
| 10 % | 5 % | 0.84 | 0.86 | 0.84 | 0.84 | 0.69 | 0.83 | 15465.50 |
|  | 10 % | 0.83 | 0.87 | 0.83 | 0.84 | 0.68 | 0.82 | 15135.49 |
|  | 15 % | 0.79 | 0.87 | 0.79 | 0.82 | 0.67 | 0.82 | 14675.58 |
|  | 20 % | 0.83 | 0.85 | 0.83 | 0.83 | 0.69 | 0.83 | 14758.57 |
| 15 % | 5 % | 0.77 | 0.83 | 0.77 | 0.77 | 0.64 | 0.80 | 16573.58 |
|  | 10 % | 0.75 | 0.83 | 0.75 | 0.77 | 0.66 | 0.80 | 16472.99 |
|  | 15 % | 0.76 | 0.86 | 0.76 | 0.79 | 0.65 | 0.80 | 16799.43 |
|  | 20 % | 0.76 | 0.84 | 0.76 | 0.77 | 0.67 | 0.81 | 15495.56 |
| 20 % | 5 % | 0.77 | 0.87 | 0.77 | 0.81 | 0.67 | 0.81 | 15932.48 |
|  | 10 % | 0.80 | 0.86 | 0.80 | 0.81 | 0.65 | 0.81 | 15823.57 |
|  | 15 % | 0.82 | 0.83 | 0.82 | 0.80 | 0.64 | 0.79 | 14597.92 |
|  | 20 % | 0.73 | 0.82 | 0.73 | 0.74 | 0.63 | 0.79 | 8885.70 |

**Table 5:** Robustness results for the whole test set (random error seq.) for Keras Classifier with *k*-mers and different % of errors.

| % of Seq. | % of Error in each Seq. | Acc. | Prec. | Recall | F1 (weighted) | F1 (Macro) | ROC-AUC | Train. runtime (sec.) |
| --- | --- | --- | --- | --- | --- | --- | --- | --- |
| 5 % | 5 % | 0.18 | 0.64 | 0.18 | 0.16 | 0.10 | 0.55 | 11832.46 |
|  | 10 % | 0.08 | 0.65 | 0.08 | 0.06 | 0.02 | 0.51 | 9405.35 |
|  | 15 % | 0.28 | 0.58 | 0.28 | 0.28 | 0.07 | 0.53 | 9912.31 |
|  | 20 % | 0.56 | 0.36 | 0.56 | 0.43 | 0.06 | 0.52 | 13029.71 |
| 10 % | 5 % | 0.62 | 0.59 | 0.62 | 0.54 | 0.10 | 0.55 | 13840.43 |
|  | 10 % | 0.61 | 0.55 | 0.61 | 0.52 | 0.09 | 0.54 | 14016.53 |
|  | 15 % | 0.49 | 0.54 | 0.49 | 0.49 | 0.07 | 0.54 | 14038.03 |
|  | 20 % | 0.58 | 0.45 | 0.58 | 0.50 | 0.09 | 0.54 | 14202.82 |
| 15 % | 5 % | 0.27 | 0.65 | 0.27 | 0.26 | 0.06 | 0.54 | 14790.52 |
|  | 10 % | 0.16 | 0.10 | 0.16 | 0.07 | 0.05 | 0.52 | 14539.64 |
|  | 15 % | 0.19 | 0.56 | 0.19 | 0.18 | 0.04 | 0.52 | 13956.71 |
|  | 20 % | 0.04 | 0.48 | 0.04 | 0.03 | 0.01 | 0.50 | 13321.69 |
| 20 % | 5 % | 0.60 | 0.71 | 0.60 | 0.58 | 0.14 | 0.57 | 14172.11 |
|  | 10 % | 0.22 | 0.58 | 0.22 | 0.19 | 0.08 | 0.53 | 12912.32 |
|  | 15 % | 0.46 | 0.57 | 0.46 | 0.43 | 0.05 | 0.52 | 8594.60 |
|  | 20 % | 0.03 | 0.59 | 0.03 | 0.01 | 0.01 | 0.50 | 13884.36 |

To visually compare the accuracy of the average accuracy for the consecutive errored sequences approach, we plot the average accuracies in Figure 1a. Similarly, Figure 2 contains the robustness results for the same two approaches.

**Fig. 1:**
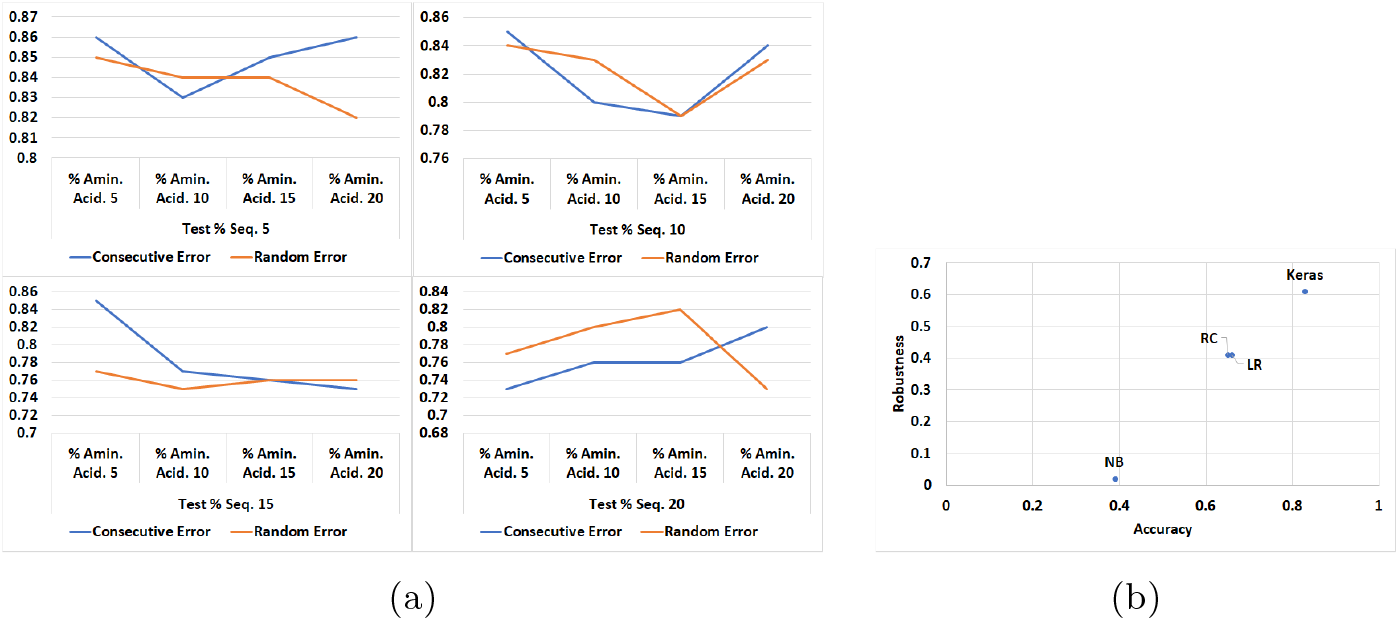
(a)Accuracy Comparison of the Consecutive error generation and Random error generation approaches for different fractions of adversarial spike sequences. (b) Accuracy (x-axis) vs Robustness (y-axis) plot (for average accuracy values) for different ML and DL methods for 10% adversarial sequences from the test set with 10% amino acids flips.

**Fig. 2:**
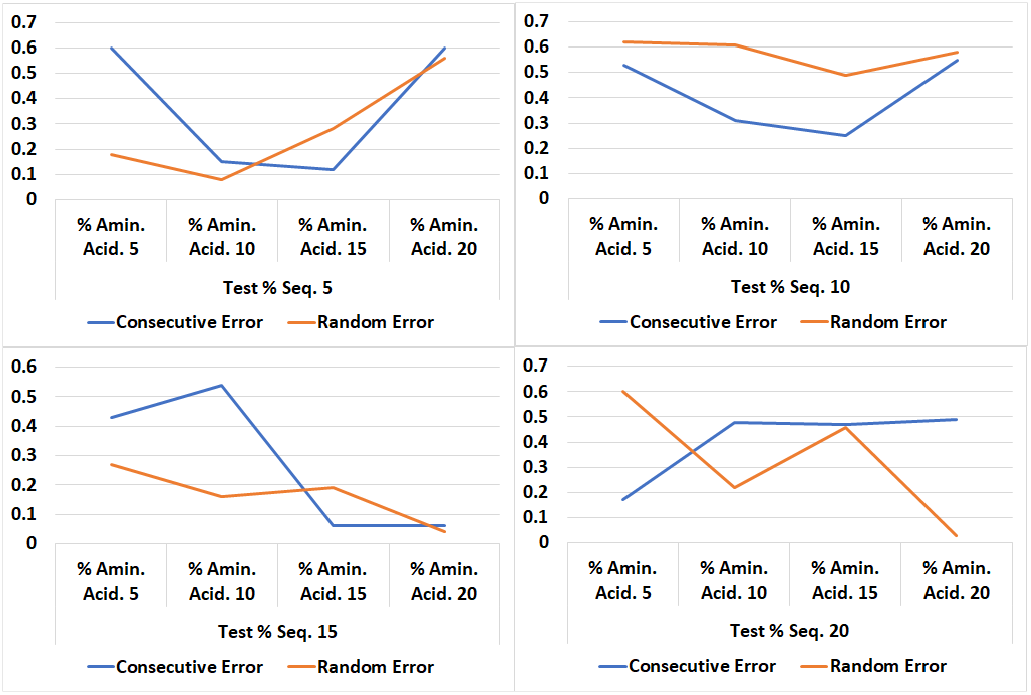
Robustness Comparison of the Consecutive error generation and Random error generation approaches for different fractions of adversarial spike sequences.

The accuracy and the robustness results for the other ML models (using the Spike2Vec approach) are given in Tables 6 and Table 7, respectively. From the results, we can conclude that our deep learning-based model is more accurate and robust than other compared machine learning models. Another interesting outcome from the results is that the *k*-mers-based feature vector is more robust than traditional one-hot embedding. This is a kind of “proof of concept” that since *k*-mers preserve the order of the amino acids in a spike sequence (as order matters in genomic sequence data), it outperforms the traditional OHE by a significant margin.

**Table 6:**
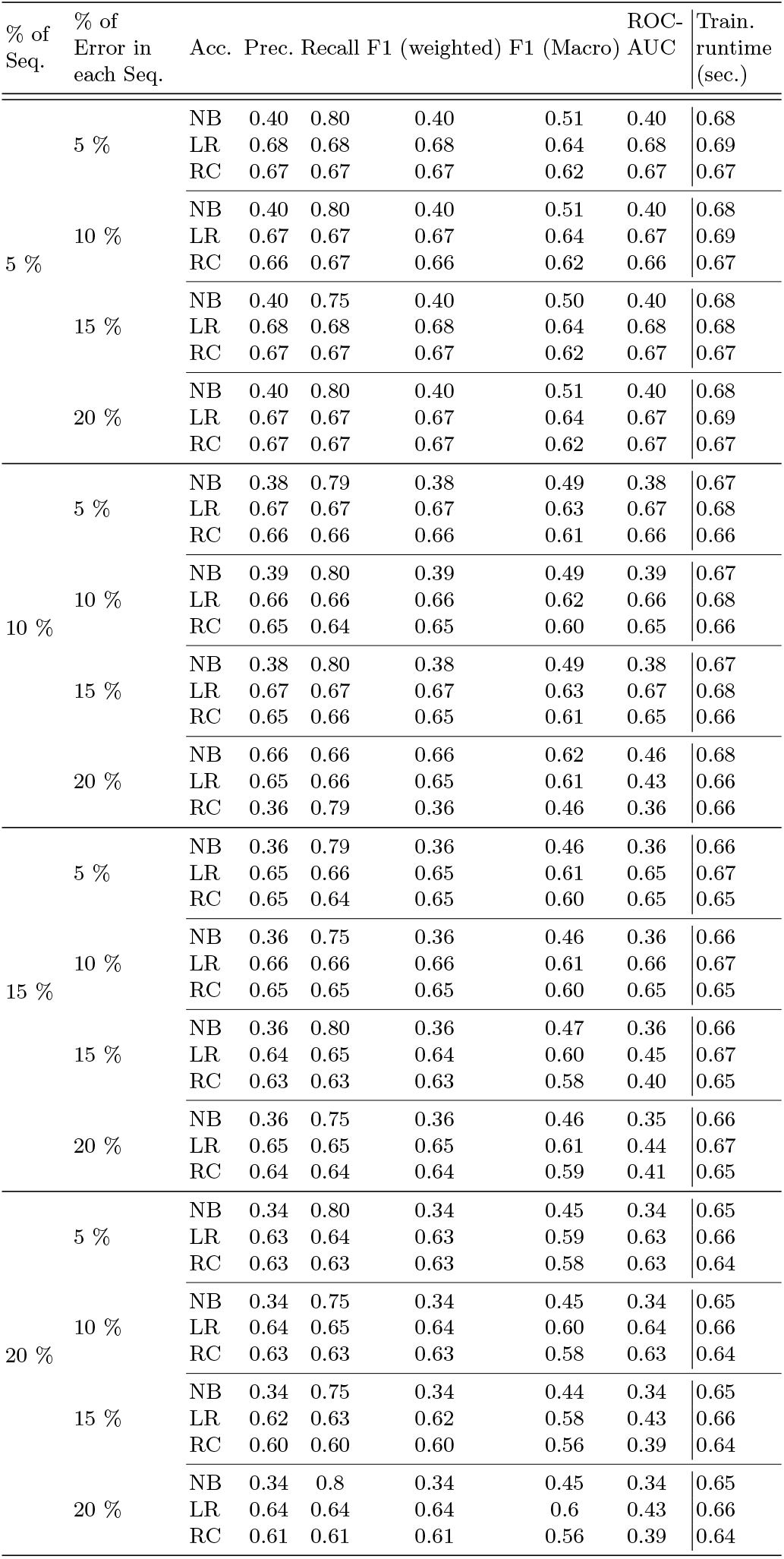
Accuracy results for the whole test set (random error seq.) for ML models with Spike2Vec approach and different % of errors.

| % of Seq. | % of Error in each Seq. | Acc. | Prec. | Recall | F1 (weighted) | F1 (Macro) | ROC-AUC | Train. runtime (sec.) |
| --- | --- | --- | --- | --- | --- | --- | --- | --- |
| 5 % | 5 % | NB | 0.40 | 0.80 | 0.40 | 0.51 | 0.40 | 0.68 |
|  |  | LR | 0.68 | 0.68 | 0.68 | 0.64 | 0.68 | 0.69 |
|  |  | RC | 0.67 | 0.67 | 0.67 | 0.62 | 0.67 | 0.67 |
|  | 10 % | NB | 0.40 | 0.80 | 0.40 | 0.51 | 0.40 | 0.68 |
|  |  | LR | 0.67 | 0.67 | 0.67 | 0.64 | 0.67 | 0.69 |
|  |  | RC | 0.66 | 0.67 | 0.66 | 0.62 | 0.66 | 0.67 |
|  | 15 % | NB | 0.40 | 0.75 | 0.40 | 0.50 | 0.40 | 0.68 |
|  |  | LR | 0.68 | 0.68 | 0.68 | 0.64 | 0.68 | 0.68 |
|  |  | RC | 0.67 | 0.67 | 0.67 | 0.62 | 0.67 | 0.67 |
|  | 20 % | NB | 0.40 | 0.80 | 0.40 | 0.51 | 0.40 | 0.68 |
|  |  | LR | 0.67 | 0.67 | 0.67 | 0.64 | 0.67 | 0.69 |
|  |  | RC | 0.67 | 0.67 | 0.67 | 0.62 | 0.67 | 0.67 |
| 10 % | 5 % | NB | 0.38 | 0.79 | 0.38 | 0.49 | 0.38 | 0.67 |
|  |  | LR | 0.67 | 0.67 | 0.67 | 0.63 | 0.67 | 0.68 |
|  |  | RC | 0.66 | 0.66 | 0.66 | 0.61 | 0.66 | 0.66 |
|  | 10 % | NB | 0.39 | 0.80 | 0.39 | 0.49 | 0.39 | 0.67 |
|  |  | LR | 0.66 | 0.66 | 0.66 | 0.62 | 0.66 | 0.68 |
|  |  | RC | 0.65 | 0.64 | 0.65 | 0.60 | 0.65 | 0.66 |
|  | 15 % | NB | 0.38 | 0.80 | 0.38 | 0.49 | 0.38 | 0.67 |
|  |  | LR | 0.67 | 0.67 | 0.67 | 0.63 | 0.67 | 0.68 |
|  |  | RC | 0.65 | 0.66 | 0.65 | 0.61 | 0.65 | 0.66 |
|  | 20 % | NB | 0.66 | 0.66 | 0.66 | 0.62 | 0.46 | 0.68 |
|  |  | LR | 0.65 | 0.66 | 0.65 | 0.61 | 0.43 | 0.66 |
|  |  | RC | 0.36 | 0.79 | 0.36 | 0.46 | 0.36 | 0.66 |
| 15 % | 5 % | NB | 0.36 | 0.79 | 0.36 | 0.46 | 0.36 | 0.66 |
|  |  | LR | 0.65 | 0.66 | 0.65 | 0.61 | 0.65 | 0.67 |
|  |  | RC | 0.65 | 0.64 | 0.65 | 0.60 | 0.65 | 0.65 |
|  | 10 % | NB | 0.36 | 0.75 | 0.36 | 0.46 | 0.36 | 0.66 |
|  |  | LR | 0.66 | 0.66 | 0.66 | 0.61 | 0.66 | 0.67 |
|  |  | RC | 0.65 | 0.65 | 0.65 | 0.60 | 0.65 | 0.65 |
|  | 15 % | NB | 0.36 | 0.80 | 0.36 | 0.47 | 0.36 | 0.66 |
|  |  | LR | 0.64 | 0.65 | 0.64 | 0.60 | 0.45 | 0.67 |
|  |  | RC | 0.63 | 0.63 | 0.63 | 0.58 | 0.40 | 0.65 |
|  | 20 % | NB | 0.36 | 0.75 | 0.36 | 0.46 | 0.35 | 0.66 |
|  |  | LR | 0.65 | 0.65 | 0.65 | 0.61 | 0.44 | 0.67 |
|  |  | RC | 0.64 | 0.64 | 0.64 | 0.59 | 0.41 | 0.65 |
| 20 % | 5 % | NB | 0.34 | 0.80 | 0.34 | 0.45 | 0.34 | 0.65 |
|  |  | LR | 0.63 | 0.64 | 0.63 | 0.59 | 0.63 | 0.66 |
|  |  | RC | 0.63 | 0.63 | 0.63 | 0.58 | 0.63 | 0.64 |
|  | 10 % | NB | 0.34 | 0.75 | 0.34 | 0.45 | 0.34 | 0.65 |
|  |  | LR | 0.64 | 0.65 | 0.64 | 0.60 | 0.64 | 0.66 |
|  |  | RC | 0.63 | 0.63 | 0.63 | 0.58 | 0.63 | 0.64 |
|  | 15 % | NB | 0.34 | 0.75 | 0.34 | 0.44 | 0.34 | 0.65 |
|  |  | LR | 0.62 | 0.63 | 0.62 | 0.58 | 0.43 | 0.66 |
|  |  | RC | 0.60 | 0.60 | 0.60 | 0.56 | 0.39 | 0.64 |
|  | 20 % | NB | 0.34 | 0.8 | 0.34 | 0.45 | 0.34 | 0.65 |
|  |  | LR | 0.64 | 0.64 | 0.64 | 0.6 | 0.43 | 0.66 |
|  |  | RC | 0.61 | 0.61 | 0.61 | 0.56 | 0.39 | 0.64 |

**Table 7:** Robustness results for the whole test set (random error seq.) for ML models with Spike2Vec approach and different % of errors.

| % of Seq. | % of Error in each Seq. | Acc. | Prec. | Recall | F1 (weighted) | F1 (Macro) | ROC-AUC | Train. runtime (sec.) |
| --- | --- | --- | --- | --- | --- | --- | --- | --- |
| 5 % | 5 % | NB | 0.02 | 0.03 | 0.02 | 0.01 | 0.02 | 0.50 |
|  |  | LR | 0.46 | 0.29 | 0.46 | 0.35 | 0.46 | 0.50 |
|  |  | RC | 0.46 | 0.28 | 0.46 | 0.34 | 0.46 | 0.50 |
|  | 10 % | NB | 0.02 | 0.00 | 0.02 | 0.00 | 0.02 | 0.50 |
|  |  | LR | 0.41 | 0.27 | 0.41 | 0.32 | 0.41 | 0.50 |
|  |  | RC | 0.43 | 0.28 | 0.43 | 0.33 | 0.43 | 0.50 |
|  | 15 % | NB | 0.02 | 0.05 | 0.02 | 0.01 | 0.02 | 0.50 |
|  |  | LR | 0.46 | 0.28 | 0.46 | 0.34 | 0.46 | 0.50 |
|  |  | RC | 0.46 | 0.28 | 0.46 | 0.34 | 0.46 | 0.50 |
|  | 20 % | NB | 0.02 | 0.03 | 0.02 | 0.00 | 0.02 | 0.50 |
|  |  | LR | 0.41 | 0.26 | 0.41 | 0.31 | 0.41 | 0.50 |
|  |  | RC | 0.41 | 0.25 | 0.41 | 0.31 | 0.41 | 0.50 |
| 10 % | 5 % | NB | 0.02 | 0.01 | 0.02 | 0.01 | 0.02 | 0.50 |
|  |  | LR | 0.46 | 0.29 | 0.46 | 0.35 | 0.46 | 0.50 |
|  |  | RC | 0.47 | 0.30 | 0.47 | 0.36 | 0.47 | 0.50 |
|  | 10 % | NB | 0.02 | 0.00 | 0.02 | 0.00 | 0.02 | 0.50 |
|  |  | LR | 0.41 | 0.27 | 0.41 | 0.31 | 0.41 | 0.50 |
|  |  | RC | 0.41 | 0.27 | 0.41 | 0.31 | 0.41 | 0.50 |
|  | 15 % | NB | 0.02 | 0.01 | 0.02 | 0.01 | 0.02 | 0.50 |
|  |  | LR | 0.41 | 0.26 | 0.41 | 0.31 | 0.41 | 0.50 |
|  |  | RC | 0.42 | 0.26 | 0.42 | 0.31 | 0.42 | 0.50 |
|  | 20 % | NB | 0.02 | 0.01 | 0.02 | 0.01 | 0.01 | 0.5 |
|  |  | LR | 0.47 | 0.3 | 0.47 | 0.36 | 0.04 | 0.51 |
|  |  | RC | 0.48 | 0.31 | 0.48 | 0.37 | 0.05 | 0.51 |
| 15 % | 5 % | NB | 0.02 | 0.03 | 0.02 | 0.01 | 0.02 | 0.50 |
|  |  | LR | 0.46 | 0.29 | 0.46 | 0.35 | 0.46 | 0.50 |
|  |  | RC | 0.41 | 0.26 | 0.41 | 0.31 | 0.41 | 0.50 |
|  | 10 % | NB | 0.02 | 0.03 | 0.02 | 0.01 | 0.02 | 0.50 |
|  |  | LR | 0.41 | 0.27 | 0.41 | 0.32 | 0.41 | 0.50 |
|  |  | RC | 0.37 | 0.26 | 0.37 | 0.30 | 0.37 | 0.50 |
|  | 15 % | NB | 0.02 | 0.01 | 0.02 | 0.01 | 0.01 | 0.50 |
|  |  | LR | 0.48 | 0.31 | 0.48 | 0.36 | 0.05 | 0.51 |
|  |  | RC | 0.48 | 0.31 | 0.48 | 0.36 | 0.05 | 0.51 |
|  | 20 % | NB | 0.01 | 0.02 | 0.01 | 0.01 | 0.01 | 0.50 |
|  |  | LR | 0.30 | 0.22 | 0.30 | 0.25 | 0.03 | 0.49 |
|  |  | RC | 0.36 | 0.24 | 0.36 | 0.29 | 0.03 | 0.50 |
| 20 % | 5 % | NB | 0.02 | 0.03 | 0.02 | 0.01 | 0.02 | 0.50 |
|  |  | LR | 0.36 | 0.25 | 0.36 | 0.28 | 0.36 | 0.49 |
|  |  | RC | 0.36 | 0.24 | 0.36 | 0.28 | 0.36 | 0.49 |
|  | 10 % | NB | 0.02 | 0.02 | 0.02 | 0.01 | 0.02 | 0.50 |
|  |  | LR | 0.36 | 0.25 | 0.36 | 0.29 | 0.36 | 0.50 |
|  |  | RC | 0.36 | 0.26 | 0.36 | 0.28 | 0.36 | 0.49 |
|  | 15 % | NB | 0.03 | 0.03 | 0.03 | 0.01 | 0.01 | 0.50 |
|  |  | LR | 0.41 | 0.28 | 0.41 | 0.32 | 0.04 | 0.50 |
|  |  | RC | 0.46 | 0.29 | 0.46 | 0.35 | 0.04 | 0.50 |
|  | 20 % | NB | 0.02 | 0.06 | 0.02 | 0.01 | 0.01 | 0.50 |
|  |  | LR | 0.47 | 0.31 | 0.47 | 0.36 | 0.04 | 0.51 |
|  |  | RC | 0.42 | 0.29 | 0.42 | 0.33 | 0.04 | 0.50 |

The accuracy vs robustness comparison (for average accuracy values) of different ML and DL methods for 10% adversarial sequences from the test set with 10% amino acids flips is given in Figure 1b. We can see that keras classifier performs best as compared to the other ML methods. This shows that not only our DL method show better predictive performance, but is also more robust as compared to the other ML models.

## 6 Conclusion

One interesting future extension is using other alignment-free methods such as Minimizers, which have been successful in representing metagenomics data. Since an intra-host viral population can be viewed as a metagenomics sample, this could be appropriate in this context. Another future direction is introducing more adversarial attacks resembling, in more detail, the error profiles of specific sequencing technologies. One could even fine-tune this to the particular experimental setting in which one obtained a sample, similar to sequencing reads simulators such as PBSIM (for PacBio reads).

## Footnotes

3 https://drive.google.com/drive/folders/1-YmIM8ipFpj-glr9hSF3t6VuofrpgWUa?usp=sharing

4 https://www.gisaid.org/

